# ThermoBRET: a ligand-engagement nanoscale thermostability assay applied to GPCRs

**DOI:** 10.1101/2020.08.05.237982

**Authors:** Bradley L. Hoare, David N. Tippett, Amandeep Kaur, Sean A. Cullum, Tamara Miljuš, Eline J. Koers, Clare R. Harwood, Nicola Dijon, Nicholas D. Holliday, David A. Sykes, Dmitry B. Veprintsev

**Affiliations:** Centre of Membrane Proteins and Receptors (COMPARE), University of Birmingham and University of Nottingham, Midlands, UK; Division of Physiology, Pharmacology & Neuroscience, School of Life Sciences, University of Nottingham, Nottingham, NG7 2UH, UK; Institute of Metabolism and Systems Research, College of Medical and Dental Sciences, University of Birmingham, Birmingham B15 2TT, UK

## Abstract

Measurements of membrane protein thermostability allows indirect detection of ligand binding. Current thermostability assays require protein purification or rely on pre-existing radiolabelled or fluorescent ligands, limiting their application to established target proteins. Alternative methods detect protein aggregation which requires sufficiently high level of protein expression.

Here, we present a ThermoBRET method to quantify the relative thermostability of G protein coupled receptors (GPCRs), using cannabinoid receptors (CB_1_ and CB_2_) and the β_2_-adrenoceptor (β_2_AR) as model systems. ThermoBRET reports receptor unfolding, does not need labelled ligands and can be used with non-purified proteins. It uses Bioluminescence Resonance Energy Transfer (BRET) between Nanoluciferase (Nluc) and a thiol-reactive fluorescent dye that binds cysteines exposed by unfolding. We demonstrate that the melting point (T_m_) of Nluc-fused GPCRs can be determined in non-purified detergent solubilised membrane preparations or solubilised whole cells, revealing differences in thermostability for different solubilising conditions and in the presence of stabilising ligands. We extended the range of the assay by developing the thermostable tsNLuc by incorporating mutations from the fragments of split-Nluc (*T*_m_ of 87 ⁰C vs 59 ⁰C). ThermoBRET allows determination of GPCR thermostability, which is useful for protein purification optimisation and as part of drug discovery screening strategies.

## Introduction

G protein coupled receptors (GPCRs) are a large family of membrane proteins that are important drug discovery targets (Hauser, Attwood et al. 2017). Structural and biophysical studies of GPCRs have significant importance in modern drug discovery (Congreve, de Graaf et al. 2020) but one major hurdle is their successful solubilisation from their native membrane environment and subsequent purification. Optimisation of receptor stability during this process is a key component to success (Tate 2010). Additionally, the ability of a bound ligand to stabilise the receptor structure is a property which can be exploited in screening efforts to find novel drug candidates (Fang 2012, Zhang, Stevens et al. 2015).

Existing GPCR protein stability assays rely on the availability of a high-affinity radioligand to act as a tracer for receptor functionality (Galvez, Parmentier et al. 1999, Serrano-Vega, Magnani et al. 2008, Robertson, Jazayeri et al. 2011, Magnani, Serrano-Vega et al. 2016). In the absence of the radioactive tracer, temperature-induced aggregation-based techniques such as technology developed by Heptares Therapeutics (now Sosei Heptares) (Marshall, Jazayeri et al. 2013) or temperature shift fluorescence size exclusion chromatography TS-FSEC (Hattori, Hibbs et al. 2012, Vuckovic 2017, Nji, Chatzikyriakidou et al. 2018) can be used. Alternative fluorescence-based techniques with higher throughput exist, such as the N-[4-(7- diethylamino-4-methyl-3-coumarinyl)phenyl]maleimide (CPM) assay, which utilises a thiol-reactive fluorescent fluorochrome. This dye reacts with exposed cysteines, acting as a sensor of protein stability in the temperature-dependent unfolding process (Alexandrov, Mileni et al. 2008). Other thiol-reactive dyes such as BODIPY-FL-Cystine (BLC) or 4-(aminosulfonyl)-7-fluoro-2,1,3-benzoxadiazole (ABD) are also available for stability measurements (Isom, Marguet et al. 2011, Bergsdorf, Fiez-Vandal et al. 2016). However, both these techniques currently require purified protein in microgram quantities which is a considerable drawback.

Low abundance of GPCRs even in over-expressing systems and their inherently low stability in detergents (Milic and Veprintsev 2015) calls for sensitive protein stability assays that can be used without protein purification or pre-existing tracer compounds.

Here, we present the ThermoBRET assay based on bioluminescence resonance energy transfer between the bright Nanoluciferase (Nluc, and, correspondingly, NanoBRET) (Hall, Unch et al. 2012), acting as a donor of light and a thiol reactive Sulfo-Cyanine3 maleimide (SCM) dye, the acceptor, allowing us to quantify the relative thermostability of non-purified GPCRs solubilised into detergent micelles. As a test case we focus on two GPCRs, the cannabinoid receptor 2 (CB_2_) as a therapeutically promising (Pertwee 2012) but unstable drug target (Vukoti, Kimura et al. 2012, Beckner, Gawrisch et al. 2019, Beckner, Zoubak et al. 2020) and the previously well characterised β_2_ adrenergic (β_2_AR) receptor. This assay detects picomolar concentration, corresponding to nanogram amounts, of target protein. Due to the nature of the homogeneous assay format, negating the need to separate bound and unbound ligand, can be used to detect binding of low-affinity ligands. Since we employ NanoBRET detection, these assays are safer than radiometric alternatives and can be readily performed in 96- and 384-well assay format.

## Results

### ThermoBRET provides reliable measurements of GPCR stability

We fused a Nluc (Hall, Unch et al. 2012) to the receptor N-terminus, preceded by a cleaved signal peptide to ensure its’ successful expression and plasma membrane trafficking (Supplementary Information 1). Detergent solubilised receptor samples containing a thiol reactive Sulfo-Cyanine3 maleimide (SCM) acceptor are incubated at varying temperatures using a gradient forming PCR thermocycler. As the receptor unfolds on heating the SCM covalently binds to exposed cysteine residues (Figure 1). We chose SCM because of its suitability as a BRET acceptor for Nluc, water solubility, and relatively low cost compared to other thiol-reactive fluorophores. In principle, any maleimide or other thiol-reactive conjugated fluorescent dye with overlapping donor-acceptor emission-absorption spectra can be used. The unfolded state of the receptor due to thermal denaturation is measured as NanoBRET between the Nluc tag and the SCM acceptor and is quantified as a ratio of the donor and acceptor light emissions, termed the NanoBRET ratio. The relative thermostability of a receptor in different solubilised non-purified membrane preparations can be easily determined, first by thermal denaturation across a temperature gradient on a thermocycler block, rapid cooling to 4 °C, and then following the addition of the Nluc substrate furimazine and measurement of the NanoBRET ratio in a 384-well luminescence plate reader at room temperature (Figure 1). The midpoint of the transition curve is found by fitting the data to a Boltzmann sigmoidal equation to obtain a *T*_m_.

**Figure 1:**
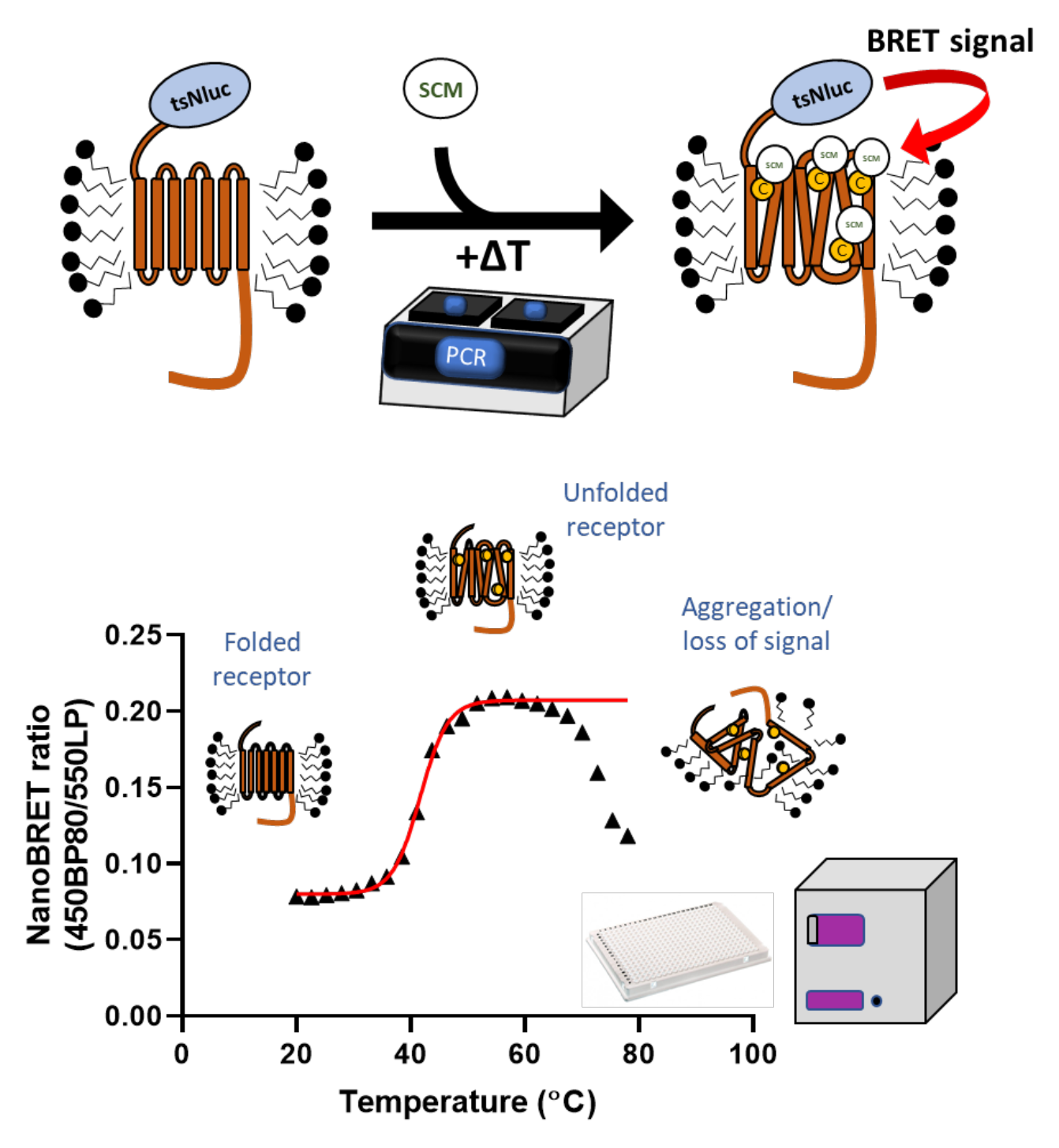
Principle of ThermoBRET assay measured in 384-well plate format. Detergent solubilised non-purified membrane preparations expressing GPCRs fused at the N-terminus with Nluc (or tsNluc) are heated using a PCR thermocycler in the presence of sulfo-Cy3 maleimide (SCM). As the protein unfolds due to thermal denaturation, SCM reacts with newly exposed cysteine residues putting the sulfo-Cy3 acceptor fluorophore in proximity with the Nluc donor. At higher temperatures, protein aggregation leads to a decrease in the NanoBRET signal and these points are truncated before fitting to a Boltzmann sigmoidal equation to obtain a melting point (*T*_m_).

### Detergents affect stability of receptors

When solubilised in DDM detergent, CB_2_ had a *T*_m_ of around 33 °C (Figure 2A) and was marginally more stable in LMNG (*T*_m_ = 35 °C). Addition of CHAPSO and the cholesterol derivative CHS in the detergent micelles provided the highest thermostability for CB_2_ (*T*_m_ = 43 °C in LMNG/CHAPSO/CHS). This observation is consistent with the reported increase in CB_2_ stability in DDM/CHAPSO mixed micelles (Vukoti, Kimura et al. 2012).

**Figure 2:**
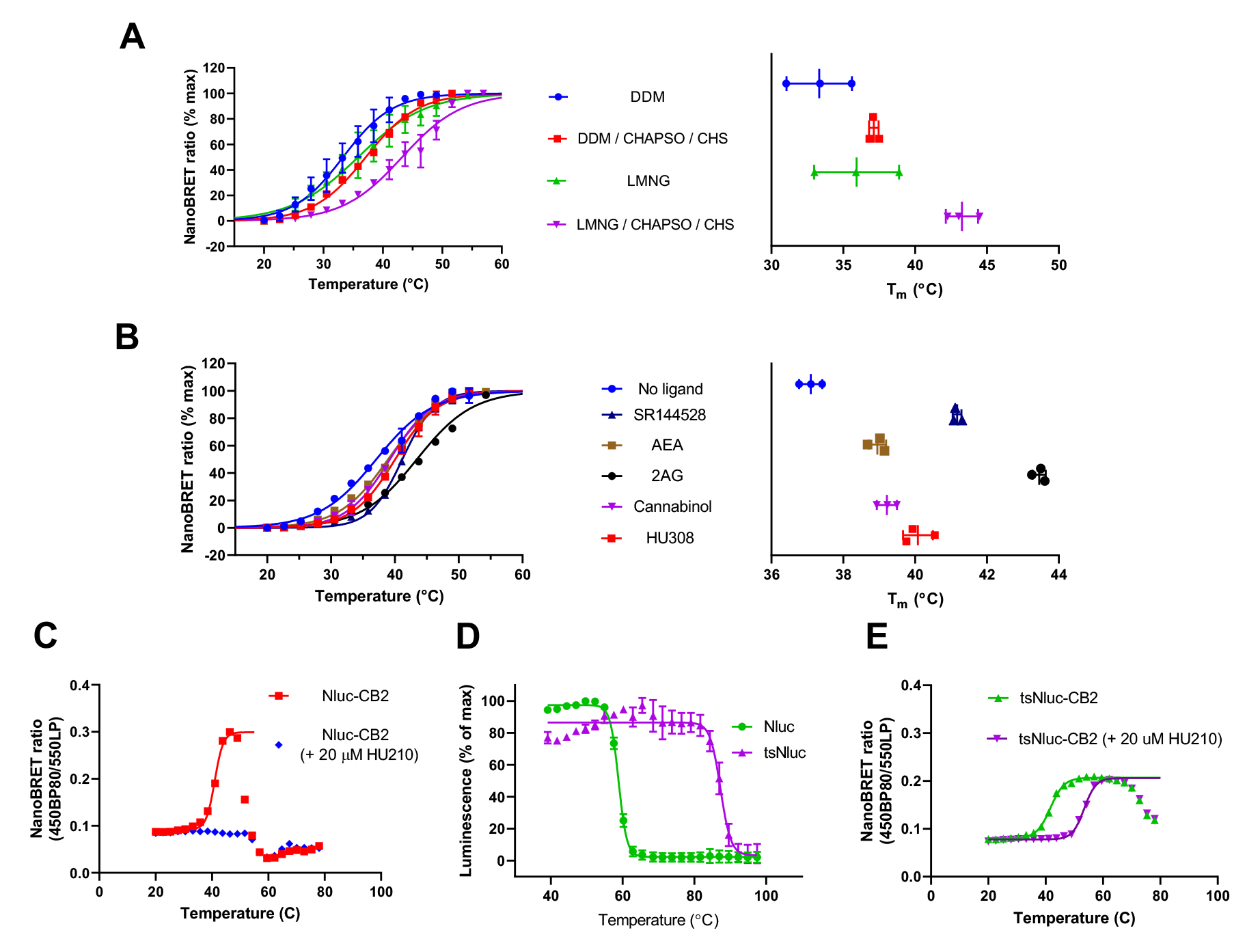
ThermoBRET measurements in different detergent conditions and with stabilising ligands, demonstrating superior performance of tsNluc over Nluc for high thermostability situations. ThermoBRET thermostability curves and pooled *T*_m_ measurements for **(A)** Nluc-CB_2_ solubilised in the indicated detergent conditions. **(B)** in DDM/CHAPSO/CHS, in the presence/absence of ligands. **(C)** ThermoBRET curve for Nluc-CB_2_ solubilised in DDM/CHAPSO/CHS, showing that the curve for the receptor bound to HU210 cannot be fitted as it is stable beyond the point of Nluc stability. **(D)** Luminescence thermostability curves of purified Nluc and tsNluc. **(E)** ThermoBRET using tsNluc-CB_2_ in the presence/absence of HU210, showing a full fit for both curves. **(A)** and **(B)** show pooled normalised data showing mean ± standard deviation for the number of experimental replicates evident in the far-right graph (n≥2). **(C)** and **(E)** are raw fitted data from a single experiment performed 3 times. **(D)** is pooled normalised data from 3-independent experiments.

Differences in the detergent stability of the adrenergic β_2_-receptor were also found (Supplementary Figure 1). Hence, this assay can be readily used to screen for the best detergent solubilising conditions before attempting a large-scale purification.

### Ligands stabilise CB_2_

We also tested a selection of endogenous and synthetic cannabinoid receptor ligands for their ability to increase the thermostability of CB_2_ (Figure 2B). The lipophilicity of its ligands has made the CB_2_ receptor a particularly challenging target for ligand binding experiments due to their high non-specific binding. These ligands were all tested at a concentration of 20 µM, well above their dissociation constant (*K*_D_) at room temperature, in order to ensure full occupancy of the solubilised receptors. Interestingly, the endogenous cannabinoid 2-arachidonoylglycerol (2AG) increased the *T*_m_ of CB_2_ by around 6 °C, whereas the other endogenous cannabinoid anandamide (AEA) only increased the *T*_m_ by around 2 °C. The most probable reason for these observations are the variable temperature dependence of the affinity of the ligand to the receptor as well as well as the degree of the entropy contribution to the binding (Layton and Hellinga 2010). Other synthetic cannabinoid ligands HU308 and SR144528 also produced appreciable increases in thermostability, and the pattern of ligand stabilisation appeared different for the related CB_1_ receptor (Supplementary Figure 2).

### tsNluc extends the range of the ThermoBRET assay

One problematic aspect of the ThermoBRET is the thermostability of the Nluc donor itself, which has been reported to unfold at around 55 - 60 °C (Hall, Unch et al. 2012). This limits the thermal range for this assay and prevents accurate *T*_m_ determination in conditions where the receptor itself is particularly thermostable, for example when CB_2_ is bound to the high affinity non-selective cannabinoid agonist HU210 (Figure 2C). We therefore combined Nluc mutations which had been developed by Promega as part of their efforts to create a stable split-luciferase system (Dixon, Schwinn et al. 2016) and found that these mutations improved thermostability of the full length luciferase by about 30 °C (Figure 2D). In line with previous reports (Hall, Unch et al. 2012) we found that purified Nluc had a *T*_m_ of 59 °C, and that purified thermostable Nluc (tsNluc) had a *T*_m_ of 87 °C (Figure 2D), making it preferable for thermostability measurements across a wide temperature range. Importantly, tsNluc contains no cysteine residues (Supplementary Information 2) and thus is unaffected by maleimide/thiol chemistry. Further characterisation showed tsNluc to have a similar luminescence emission profile as Nluc with furimazine as a substrate, although with a lower luminescence output (Supplementary Figure 3). Applying this novel tsNluc fusion improved the working temperature range of the ThermoBRET assay and allowed successful *T*_m_ determination for CB_2_ in the presence of HU210 (Figure 2E). Strikingly, HU210 was able to stabilise CB_2_ by around 12 °C, the highest level achieved of any of the CB_2_ ligands tested.

To further assess the ability of this improved assay format to determine the stability of GPCRs in detergent we created a tsNluc-β_2_AR expression construct. Previous work employing the CPM assay indicates that this receptor can be stabilised by high affinity antagonists (Wacker, Fenalti et al. 2010). Due to the higher throughput achievable with our BRET-based system we were able to assess the receptor stabilising effects of both β-adrenergic agonists and antagonists (Figure 3 A and B). Both high affinity antagonist and agonist were able to stabilise the receptor to a degree which was dependent on the affinity of the ligands for the receptor (Figure 3C). In contrast *k*_off_ and *k*_on_ were by themselves more poorly correlated with receptor stabilisation (Figure 3D and E).

**Figure 3:**
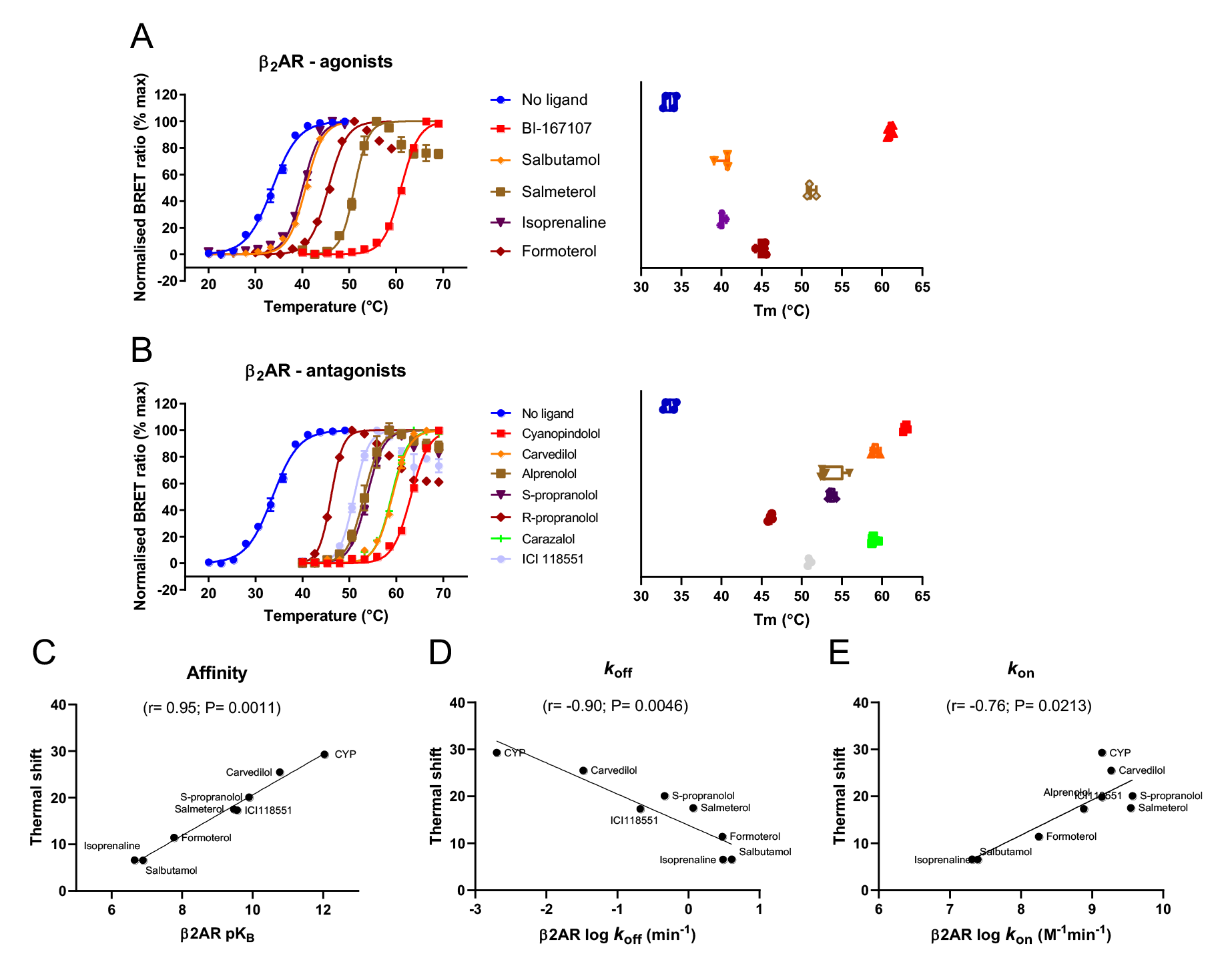
tsNluc β_2_AR thermoBRET measurements in 0.1% DDM with stabilising ligands and high thermostability situation. ThermoBRET thermostability curves and pooled *T*_m_ measurements for tsNluc-β_2_AR solubilised in DDM in the presence/absence of **(A)** agonist ligands and **(B)** antagonist ligands. The magnitude of shifts are shown on panels on the right, with ligands as indicated on the left panel. Correlations of ligand thermostability (Δ*T*_m_) with **(C)** ligand affinity, **(D)** ligand *k*_off_ and **(E)** ligand *k*_on_. **(A)** and **(B)** show pooled normalised data showing mean ± standard deviation for the number of experimental replicates evident in the far-right graph (n≥3), pooled normalised data from 3 or more independent experiments. Radioligand binding data values were taken from (Sykes, Parry et al. 2014).

### Lipid concentration in lipid-detergent micelles affects stability of the receptor

Having established a robust assay to measure receptor stability, we examined the more subtle effects of lipid-detergent ratio in the solubilised tsNluc-CB_2_. Firstly, we examined if the stability of CB_2_ is affected by the receptor concentration, while keeping the membrane fraction/detergent ratio the same by supplementing the fraction of CB_2_ membranes with “empty” non-transfected HEK293 membranes to the same total amount (Supplementary Figure 4A). The reported value was the same, suggesting that CB_2_ stability is not affected by its concentration, at least within the tested range. Secondly, we examined if an increase in membrane/detergent ratio could affect CB_2_ stability, by supplementing a fixed amount of CB_2_ membranes with an increasing amount of “empty” membranes (Supplementary Figure 4B). Surprisingly, the receptor stability was decreased by 2-3 °C at higher membrane/detergent ratio, although it plateaued at concentrations above 20 ng/μL, as generally increasing concentration of lipids may lead to stabilisation of receptors (Cecchetti, Strauss et al. 2021). A possible explanation for this is that the increased total protein concentration (HEK293 membranes contain a significant amount of protein) may accelerate aggregation process. This observation suggests that for each new target this relationship needs to be explored, and concentration of membrane and detergent should be kept constant and an appropriate point on the plateau region should be chosen.

### Stability of the receptor measured by ThermoBRET reflects a loss of ligand binding

Receptor denaturation is a complex process, progressing through a loss of tertiary structure through potential intermediates that may have the overall organisation of the correctly folded receptor (and protecting cysteines from modification) to complete loss of tertiary structure and aggregation. It is important to understand what process is sensed by the ThermoBRET assay.

Stabilisation of a receptor can also be monitored through addition of a fluorescent ligand to a detergent solubilized receptor, measured using NanoBRET detection. In this case unfolding of a target protein in response to an increase in temperature will result in a loss in specific binding. Figure 4A shows the saturation binding curve for propranolol-green binding to the tsNluc-β_2_AR receptor solubilised in DDM. The *K*_d_ of propranolol-green for the β_2_AR was determined to be 5.7 ± 1.1 nM. Figure 4B shows the loss in propranolol-green specific binding signal as the protein unfolds in response to an increase in temperature, following preincubation of a saturating concentration of fluorescent ligand (1 μM) with the receptor.

**Figure 4:**
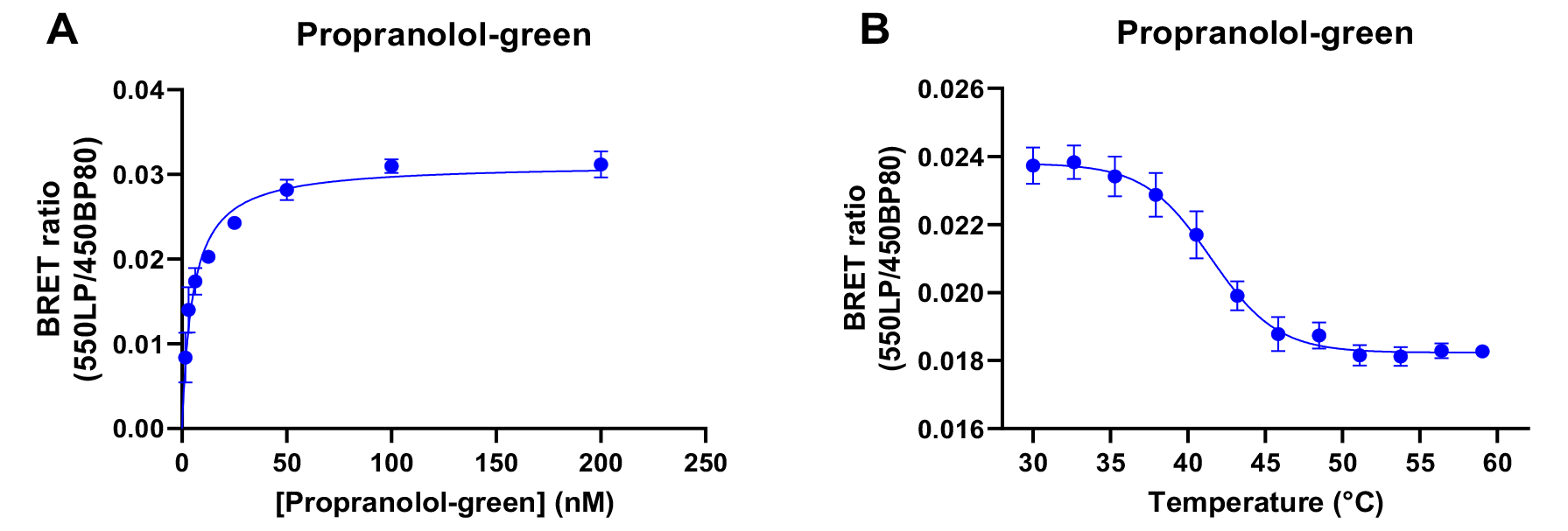
BRET based fluorescent tracer binding to the β_2_AR solubilised in DDM. (A) Saturation binding curve of propranolol-green binding the β_2_AR. ThermoBRET thermostability curve for tsNluc-β_2_AR in the presence of (1 μM) propranolol-green. Data is plotted as the mean ± standard error for 3 replicates.

### Isothermal ThermoBRET

Finally, we assessed the dependency of agonist and antagonist ligand concentration on tsNluc-β_2_AR thermostability at a constant temperature 35°C, just above the *T*_m_ of the apo form of the receptor. Receptor stability measurements expressed as a function of ligand concentration are shown in Figure 5A. ThermoBRET IC_50_ values were derived for this smaller test set and correlated with the thermal shift values obtained at fixed concentrations of individual agonist and antagonists. A linear relationship was observed between these two measures (Figure 5B). Finally, the ThermoBRET IC_50_ values were correlated with radioligand binding affinity values obtained for the different ligands (Figure 5C). Again, an excellent correlation was observed between the two data sets apart from for the very slowly dissociating antagonist cyanopindolol. There are two plausible explanations for this: either the very slow off-rate of this ligand means that for this ligand equilibrium is not achieved prior to the determination of ThermoBRET IC_50_ values, samples being kept on ice prior to melting, or we are observing the phenomenon of ligand depletion due to nM concentrations of receptor present in the reaction mixture.

**Figure 5:**
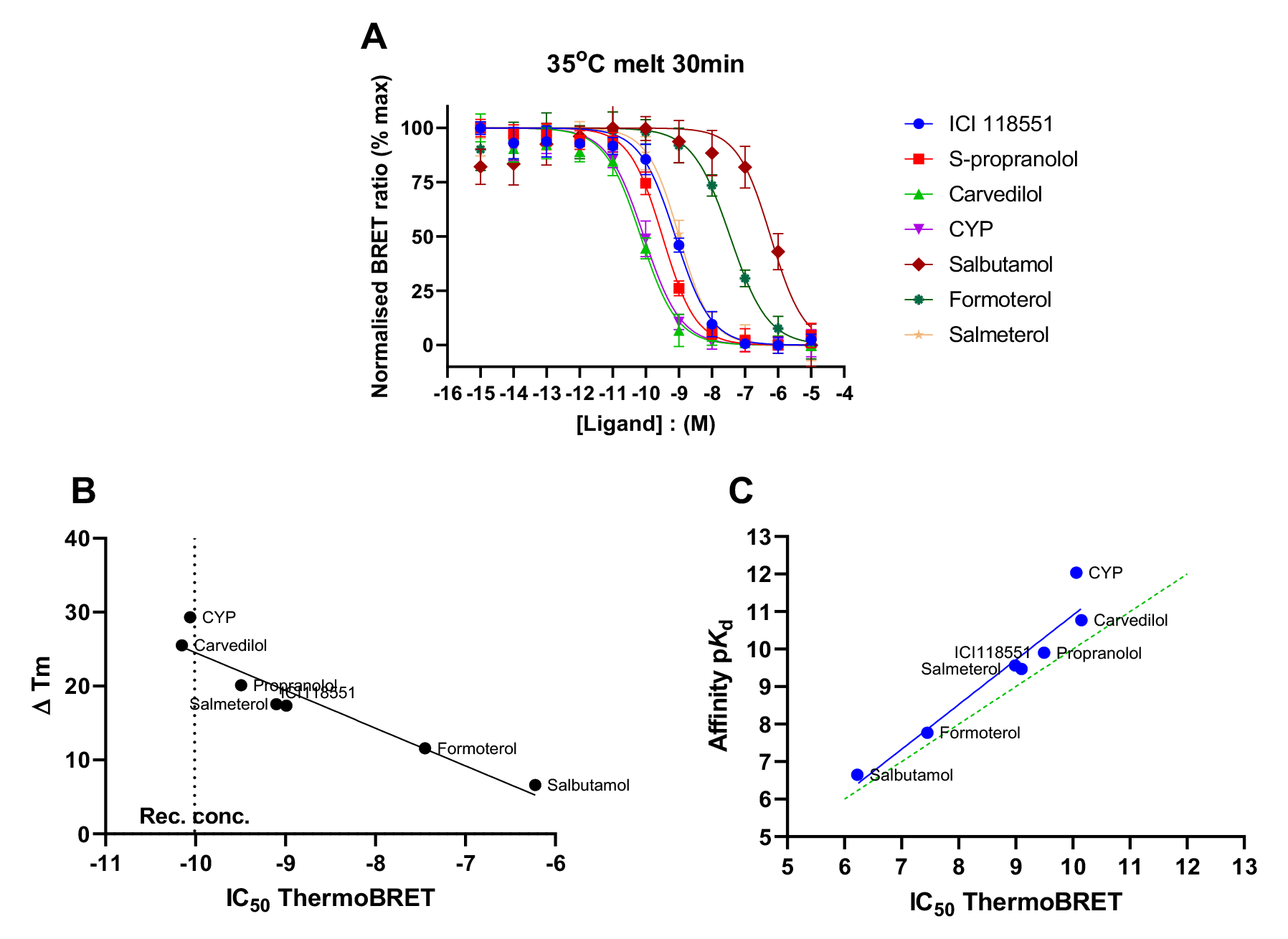
tsNluc β_2_AR thermoBRET measurements in 0.1% DDM with stabilising ligands expressed as a function of concentration. **(A)** ThermoBRET IC_50_ curves obtained at a fixed temperature of 35°C following a 30min incubation with either agonists or antagonist ligands. Normalised data is plotted as the mean ± standard deviation for 3 replicates. ThermoBRET IC_50_ values were derived for this smaller test set and correlated with **(B)** the change in thermal shift obtained at fixed concentrations of individual agonist or antagonist and **(C)** radioligand binding derived ligand affinity values. All correlations are derived from mean thermostability measurements consisting of at least 3 replicates. Radioligand binding data values were taken from (Sykes, Parry et al. 2014).

### Assay sensitivity

Solubilised receptor concentration was estimated by extrapolation from a standard curve of the luminescence emission from purified tsNluc protein of known concentrations. ThermoBRET assay sensitivity was then determined by dilution of this solubilized β_2_AR receptor sample with receptor stabilisation assessed by monitoring the change in BRET ratio as a function of increasing temperature, see Supplementary Figure 5. This confirms that ThermoBRET is a nanoscale system with sensitivity in line with that of a traditional ligand binding assay. In further experiments we tested the effect of SCM dye concentration on the thermal unfolding of the β_2_AR (see Supplementary Figure 6), confirming that dye concentrations in the region of 1-3 μM are sufficient to monitor the unfolding of GPCRs and without significant quenching of the luminescent signal.

### Receptor solubilisation directly from cells

In addition, we assessed the stability of the β_2_AR solubilised directly from whole cells and the ability of orthosteric ligands to stabilise the receptor. Figure 6 shows the measured stability of the apo β_2_AR solubilised from whole cells in the absence and presence of a selection of antagonist and agonist ligands and the resulting correlation of *T*_m_ values obtained with the same ligands from membrane solubilised receptors. The measurement of receptor thermostability directly from whole cells further reduces the number of steps required for the assay (ie. no need to prepare membranes) and allows target engagement to be assayed when receptors are still in a native membrane environment if it is added prior to solubilisation.

**Figure 6:**
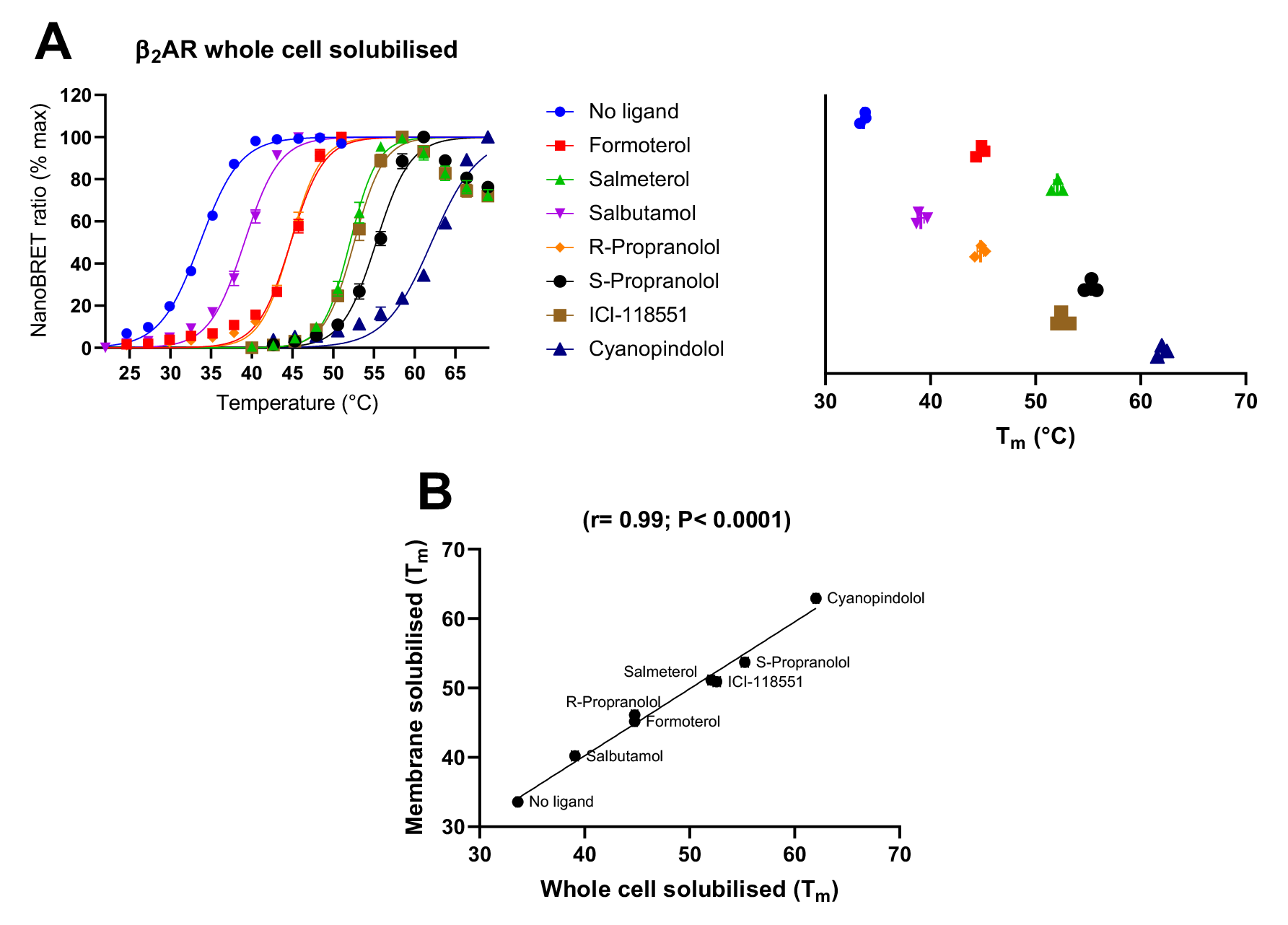
tsNluc β_2_AR thermoBRET measurements in 0.1% DDM following solubilisation from whole cells. ThermoBRET thermostability curves and pooled *T*_m_ measurements for tsNluc-β_2_AR solubilised in DDM in the presence/absence of **(A)** agonist ligands and antagonist ligands. The magnitude of shifts is shown on the panel on the right **(B)** shows the correlation between *T*_m_’s determined following solubilisation of the β_2_AR from HEK293TR membranes or whole cells. **(A)** and **(B)** show pooled normalised data showing mean ± standard deviation for the number of experimental replicates evident in the far-right graph (n≥3), pooled normalised data from 3 or more independent experiments.

## Discussion

Here, we aimed to establish sensitive technique that allows to measure receptor stability in crude detergent-solubilised preparations, without a need for a unique tracer compound. ThermoBRET proved to be sensitive and selectively reporting protein stability of solubilised membrane preparations and even directly solubilised cells. It utilises universal cysteine-reactive fluorescent dyes for readout and can be used to detect binding of specific non-fluorescent ligands.

The processes of protein unfolding and protein aggregation are related, yet very separate, phenomena. As mentioned in the introduction, a decrease in GFP fluorescence (Vuckovic 2017, Nji, Chatzikyriakidou et al. 2018) or luciferase signal (Marshall, Jazayeri et al. 2013) has been used for measuring temperature-induced aggregation of GPCRs. Nluc has also been successfully used in similar applications which monitor protein aggregation of soluble proteins. Such applications fuse either full length Nluc (Dart, Machleidt et al. 2018) or split Nluc (Martinez, Asawa et al. 2018) to the protein of interest and monitor the decrease in luminescence activity to measure aggregation of the protein of interest after thermal denaturation. In contrast, the ThermoBRET assay described here captures the initial conformational unfolding events which expose maleimide reactive cysteine residues in the protein of interest. In addition, ThermoBRET measurement is buffered from changes in the concentration of the luciferase fused target protein because a ratiometric method is used to calculate resonance energy transfer, contrasting with assays which measure luminescence intensity only. This means the measurement remains robust even at low concentrations of the target protein where the magnitude of the measured signal is at the lower end of detection capabilities.

We note that under the test conditions used, the luminescence activity of just 85 pM purified Nluc and tsNluc was measurable in a 96-well plate (Figure 2D), making the assay extremely sensitive. This makes the assay particularly amenable situations where there are limitations on the amounts of reagent that can be provided, for example protein targets which are poorly expressed *in vitro* and/or *in vivo*. This assay principle could even be applied in more physiologically relevant *in vivo* cellular models whereby tsNluc (or the 11 amino acid HiBiT tag) is fused to an endogenously expressed protein via CRISPR-mediated insertion (White, Johnstone et al. 2019).

The novel thermostable tsNluc we describe has clear advantages compared to Nluc. Firstly, due to its improved thermostability it is less likely to unfold before the protein of interest and cause sample aggregation and other possible artefacts. The T_m_ of tsNluc was 87 ˚C which may put an upper temperature limit on the ThermoBRET method, however such a high thermostability situation would be very unexpected for an integral membrane protein solubilised in detergents. Secondly, whilst previous reports (Hall, Unch et al. 2012) and our own data (Figure 2D) showed the *T*_m_ of Nluc to be 59 ⁰C, more recent use of Nluc to monitor protein aggregation show clear luminescence activity after protein samples had been heated to temperatures >60 ⁰C before cooling (Dart, Machleidt et al. 2018). We speculate that Nluc has propensity to spontaneously refold after thermal denaturation, and that the presence of detergents in the buffers of the latter report either aided Nluc refolding or delayed irreversible protein aggregation. In our ThermoBRET assays, the cysteine exposed upon Nluc unfolding would potentially react with the SCM and prevent its refolding, whereas tsNluc avoids these pitfalls. The cysteine-less sequence of tsNluc also allows easy *in vitro* chemical tagging of tsNluc-fusions. There is of interest in producing conjugates of Nluc fused to other biomolecules, though usually this involves incorporation of sequence-specific ligation motifs onto Nluc followed by enzyme-mediated ligation to the molecule of interest (Wang, Shao et al. 2017, Mie, Niimi et al. 2019, Wouters, Vugs et al. 2020). By introducing a cysteine at the C-terminus of tsNluc, any molecule could be conjugated to tsNluc by thiol-reactive chemistry and mild reaction conditions, enhancing its potential for protein engineering and enabling a wide scope for future applications.

In comparison to our previous ThermoFRET application using a terbium cryptate labelled receptor as a FRET donor (Tippett, Hoare et al. 2020), the ThermoBRET approach offers potential advantages. ThermoFRET requires cell surface labelling of the receptor-fused SNAP tag with the terbium cryptate donor molecule, adding to assay cost, but perhaps more importantly creating an extra labelling step that can be problematic if the tag is not readily exposed at the plasma membrane. In contrast, the use of a genetically encoded bioluminescent donor (ie. tsNluc or Nluc) omits this labelling step. This means that fused proteins which are poorly trafficked to the plasma membrane, are now amenable as they do not require labelling at the cell surface. Additionally, ThermoFRET requires more sophisticated detection by plate readers with time-resolved fluorescence detection capabilities, whereas BRET only requires a luminometer with filtered light detection that are more readily available in many labs. A comparison of biophysical techniques used in drug screening cascades is shown in Figure 7 along with the relative protein requirements and assay throughput potential of each technique.

**Figure 7:**
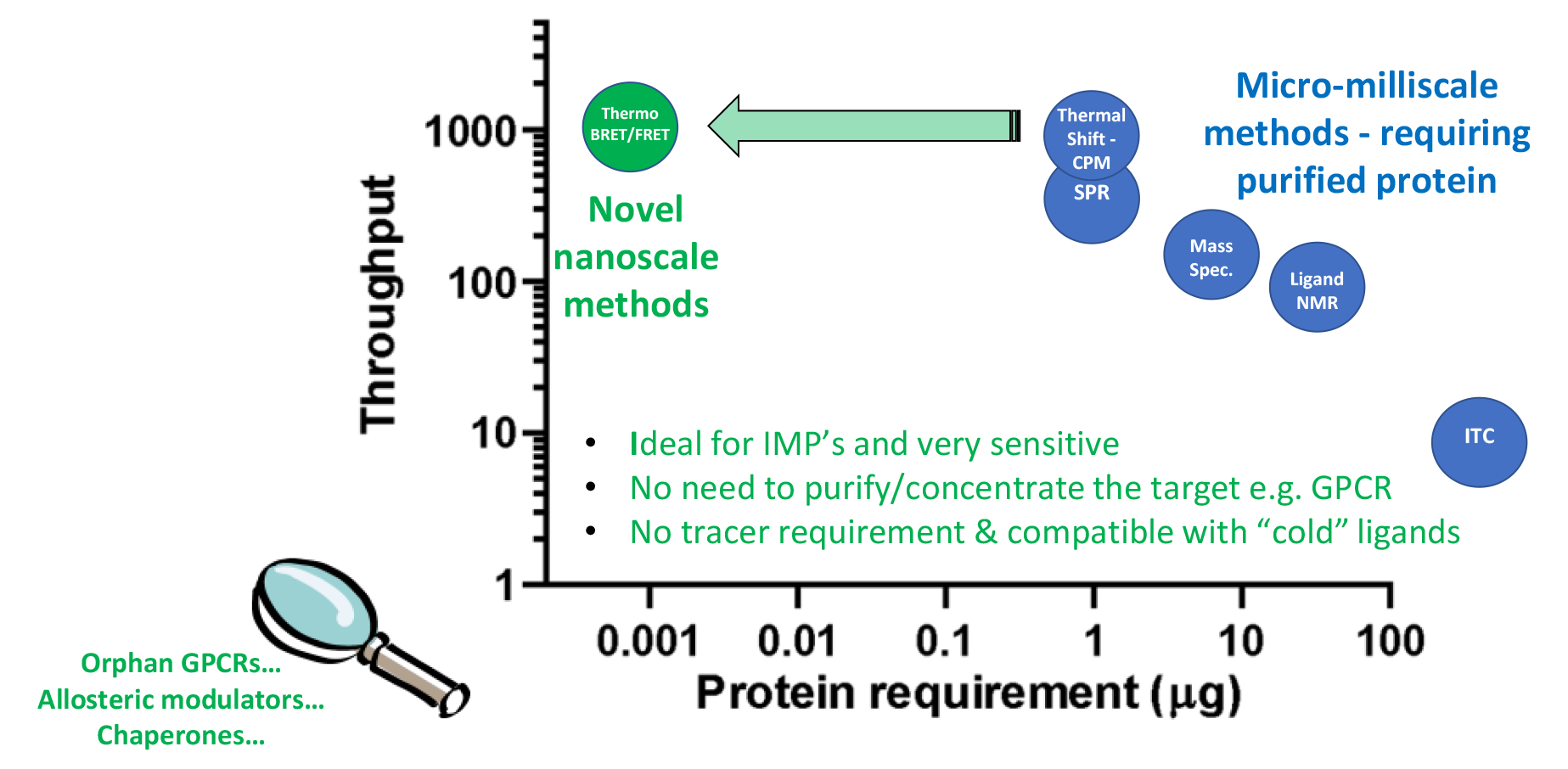
A comparison of biophysical techniques used in drug screening cascades. Outlined are the protein requirements of each technique and their estimated daily throughput screening potential, along with the advantages of ThermoBRET and some potential uses.

The ability of the ThermoBRET assay to quantify ligand-induced changes in the receptor *T*_m_ makes it an ideal tool to study ligand binding to GPCRs. *T*_m_ values obtained in the current study for the β_2_AR specific ligands compare well with those obtained previously (Zhang et al., 2015). In principle, this assay can detect compounds which bind the target at any site, assuming this interaction influences the thermodynamic conformational landscape of the protein. It can be used to screen potential ligands for orphan GPCRs as it does not depend on the availability of tool compounds or known binders to develop a competition assay. Moreover, it could be used to detect the combined stabilisation of several ligands to discover positive and negative allosteric modulators of GPCRs.

Despite a number of advantages ThermoBRET offers, and positive results obtained for β_2_AR, CB_2_ and CB_1_ receptors, this technique is not immune to the general limitations of thermal shift assays. The protein needs to be in a native state once solubilised, and some condition optimisation may need to be done for the less stable receptors. Correspondingly, it may not be successful for every target receptor tried as there is a requirement that the receptor contains buried cysteines which become exposed upon thermal denaturation. This limitation is also inherent for the CPM assay, and in situations where no free thiol exists then cysteines could be rationally introduced into the receptor sequence to generate a ThermoBRET signal.Finally, more practical experience with screening larger compound libraries will be needed to establish real life performance and limitations of this technique.

Overall, ThermoBRET is an excellent and highly sensitive tool for optimisation of solubilisation conditions and biophysical screening of GPCR compound libraries to support structural biology, aiding the drug discovery efforts.

## Methods

### Drug compounds and reagents

Sulfo-Cy3 maleimide (SCM) (Lumiprobe GmbH, Germany) was obtained in powder form, dissolved in DMSO at a concentration of 10 mM and stored in the dark at −20C. Furimazine, the substrate for Nluc, was obtained from the Nano-Glo Luciferase Assay System kit (Promega, UK) provided at a concentration of 5 mM. Cannabinoid ligands (anandamide [AEA], 2-arachydonyl-glycerol [2AG], SR144528, HU210, HU308, cannabinol) were obtained from Tocris Bioscience and dissolved in DMSO to a storage concentration of 10 mM, except AEA and 2AG which were dissolved in EtOH. Rimonabant was obtained from Roche Pharmaceuticals GmbH (Germany).

### Plasmid construction

For mammalian cell expression, receptor constructs were cloned into pcDNA4/TO using Gibson assembly (Gibson, Young et al. 2009). All GPCR constructs contained an N-terminal signal peptide (which is cleaved by signal peptidases during protein maturation and trafficking) to improve expression, followed by a TwinStrep affinity tag, then Nluc (or tsNluc) followed by the receptor sequence. The TwinStrep affinity tag was not required for these studies but was present to facilitate receptor purification and antibody-based detection if required. The synthesized cDNA for tsNluc was obtained from GeneArt Gene synthesis (Invitrogen). For bacterial cell expression of Nluc and tsNluc, cDNA sequences were cloned into the pJ411 expression plasmid with a N-terminal 10X histidine affinity tag and TEV cleavage site encoded upstream of the protein of interest. Amino acid sequences of the constructs used are provided in Supplementary Information 1. The correct sequence within the expression cassette of all plasmid constructs was verified by Sanger sequencing (Genewiz, UK).

### Mammalian cell culture

The T-Rex^TM^-293 cell line (HEK293TR; ThermoFisher Scientific) was used to make stable expressing cell lines for receptors cloned into pcDNA4/TO. HEK293TR cells were cultured in growth medium (DMEM, 10% FCS, 5 μg/mL blasticidin) in a 37 ⁰C humidified incubator with 5% CO_2_. Stable cell lines were generated by PEI transfection of pcDNA4/TO plasmids into HEK293TR cells. 24-48 hours after transfection, 20 μg/mL zeocin was incorporated into the growth medium until stable expressing, zeocin-resistant cell populations remained (2-4 weeks). To produce cells for membrane preparations, 1X T175 culture flask of confluent stable cells were treated with 1 μg/mL tetracycline for 48h to induce receptor expression. Following this, cells were lifted by trituration and centrifuged at 500 *g* for 10 minutes. Cell pellets were then frozen at −80 ⁰C until membranes were prepared.

### Membrane preparations

HEK293TR cell pellets were resuspended in 20 mL of ice cold buffer (10 mM HEPES pH 7.4, 10 mM EDTA) and homogenised using a Ultra Turrax (Ika Work GmbH, Germany). The homogenised cell suspension was then centrifuged at 4 ⁰C for 5 minutes at 500 *g* to remove whole cells and large debris, and the remaining supernatant was then centrifuged twice at 4 ⁰C and 48,000 *g* for 30 minutes before the membrane pellet was resuspended in buffer (10 mM HEPES pH 7.4, 0.1 mM EDTA). Protein concentration of resuspended membranes was determined with using Pierce BCA Protein assay kit (ThermoFisher Scientific) and was adjusted to 3 – 10 mg/mL before being aliquoted and stored at −80 ⁰C.

### ThermoBRET experiments

The CORE buffer for thermostability experiments contained 20 mM HEPES pH 7.5, 150 mM NaCl, 10% w/v glycerol, 0.5% w/v BSA. Cell membranes were diluted in CORE buffer to approximately 0.1 – 0.5 mg/mL total protein and were then centrifuged at 16,000 *g* for 60 minutes at 4 ⁰C to remove residual EDTA from the membrane preparation buffers. Membrane pellets were then resuspended in CORE buffer containing detergent, and samples were incubated at 4 ⁰C with gentle shaking for 1h to solubilise membranes. Detergent/CHS concentrations used were either 1% DDM, 1% DDM / 0.5 % CHAPSO / 0.3% CHS, 0.5% LMNG, or 0.5% LMNG / 0.5% CHAPSO / 0.3% CHS. Samples were then centrifuged again at 16,000 *g* for 60 minutes at 4 ⁰C to remove unsolubilised material, and the resulting supernatant containing detergent micelles was transferred to a fresh tube. These supernatants were then kept on ice for up to 48 hours during testing. For thermostability testing, solubilised receptors were diluted 10-fold in CORE buffer with the addition of 1 μM SCM and 20 μM of ligand (if used). This was incubated on ice for 15 minutes before being aliquoted across 96-well PCR plates and placed in the pre-cooled (4 °C) PCRmax Alpha Cycler 2 Thermal Cycler (Cole-Palmer Ltd, St. Neots, UK). Samples were then incubated at different temperatures for 30 minutes via a temperature gradient across the plate. Following rapid cooling of the samples to 4 ⁰C, samples were then transferred to white 384-well proxiplates (Perkin Elmer) containing furimazine at a final concentration of 10 μM. The plate was then read using a PHERAstar FSX plate reader (BMG) at room temperature and the 450BP80/550LP filter module. Measurements were performed in singlet for each temperature point. Whole cell experiments (10 million cells/mL of 1% DDM) were performed essentially as described above but receptor solubilisation was performed in the presence of protease inhibitor cocktail (cOmplete^TM^ mini EDTA-free Protease Inhibitor cocktail (Roche)).

### Nluc and tsNluc expression and purification

NiCo21(DE3) chemically competent E. coli were transformed with pJ411 bacterial expression plasmids and plated onto LB/agar plates containing 2% w/v glucose and 50 µg/mL kanamycin. After incubation at 37 °C for 16-24 hours, a single colony was picked to inoculate 20 mL of terrific broth containing 0.2% w/v glucose and 50 µg/mL kanamycin. After 16-24 hours in a shaking incubator set at 37 ⁰C, 15 mL of overnight culture was added to 3 L of terrific broth containing 0.2% w/v glucose and 50 µg/mL kanamycin, grown in a shaking incubator at 37 ⁰C until OD_600_ of 0.7-1, when 500 μM of isopropyl-β-D-thiogalactopyranoside (IPTG; VWR Chemicals) was added to induce protein expression. Cells were then grown overnight (16-20 hours) at 25 ⁰C in a shaking incubator before being harvested by centrifugation and frozen at −80 ⁰C. Cell pellets were then thawed on ice, and resuspended in 100 mL lysis buffer (100 mM Tris pH 7.5, 300 mM NaCl, 0.25 mg/mL chicken lysozyme, 1 µg/mL bovine DNAse I, 4 mM MgCl_2_, and 3 cOmplete^TM^ mini EDTA-free Protease Inhibitor cocktail tablets (Roche)). After 1h on ice in lysis buffer, cells were then lysed further by French press. Cell lysates were then clarified by centrifugation at 25,000 rcf for 30 minutes and then by passing through a 0.45 μm syringe filter. The His-tagged proteins from the resulting lysate were then purified using a 5mL HiTrap TALON Crude column on an ÄKTA start protein purification system (Cytiva Life Sciences) and eluted with 150 mM imidazole. Elution fractions were analysed by SDS-PAGE and fractions which contained no visible contaminants proteins were pooled together. Protein concentration was determined by A_280_ measurement on a Denovix DS-11 FX series spectrophotometer assuming the calculated molar extinction coefficient (ε_280_) of 26,930 mol^-1^.cm^-1^ for both proteins.

### Luminescence activity thermostability experiments

Purified Nluc and tsNluc proteins were serially diluted from around 200 μM down to 100 pM in CORE buffer. Proteins were then aliquoted across 96-well PCR plates (100 μL per well) and placed in the pre-cooled (4 °C) PCRmax Alpha Cycler 2 Thermal Cycler (Cole-Palmer Ltd, St. Neots, UK). Samples were then incubated at different temperatures for 30 minutes via a temperature gradient across the plate. Following rapid cooling to 4 ⁰C, 85 μL of samples were then transferred to white 96 well plates (Perkin Elmer) containing 15 μL of diluted furimazine to a final concentration of 10 μM. After 30 seconds of gentle shaking, the luminescence intensity was measured in a PHERAstar FSX plate reader at room temperature. Measurements were performed in triplicate for each temperature point, and three independent experiments were performed.

### Curve fitting and data analysis

All curve fitting and data manipulation was performed using GraphPad Prism 8. For ThermoBRET measurements, NanoBRET ratio was defined as the 550LP emission divided by the 450BP80 emission. In situations in which the NanoBRET ratio decreased at high temperatures (presumably due to protein aggregation and loss of signal), the data was manually truncated after the highest point. Data was then normalised to the upper (100%) and lower (0%) datapoints and fitted using a Boltzmann sigmoidal equation constrained to upper and lower values of 0% and 100%. For luminescence thermostability measurements, unfiltered luminescence was normalised to the top point of the dataset and fitted using a Boltzmann sigmoidal equation with no constraints.

## Data availability statement

All data used to support the conclusions are included in the paper. Raw numerical data can be obtained by contacting the corresponding authors.

## Acknowledgements

AK was funded by a COMPARE team science summer studentship (2019) awarded to BH. This research was supported by COMPARE funding to DBV. We would like to thank Uwe Grether for supplying the CB_1_ antagonist rimonabant.

---

**Supplementary Figure 1:**
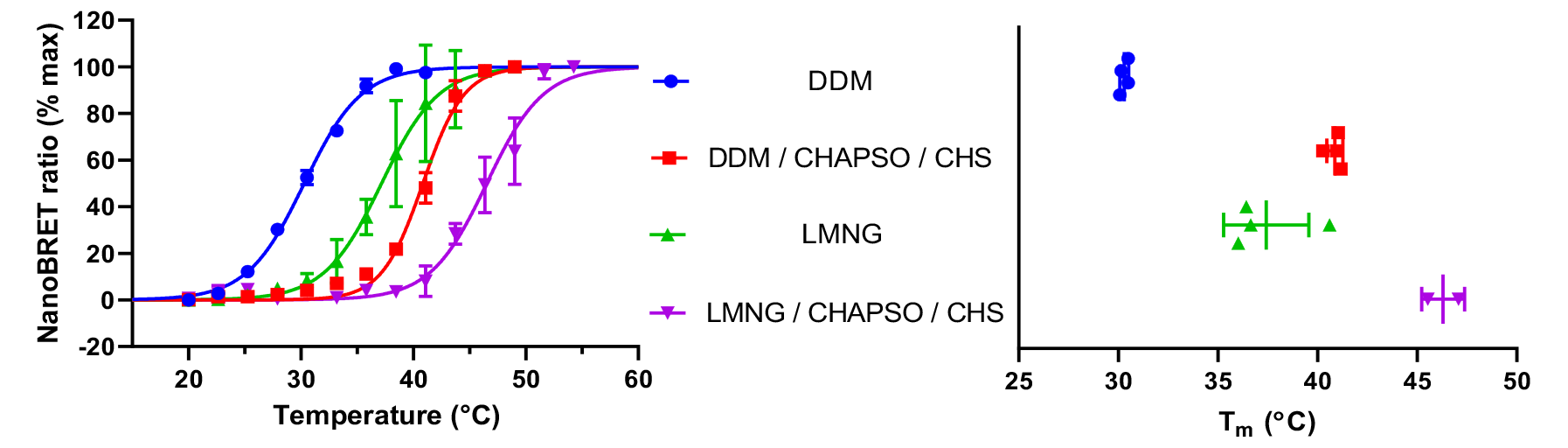
Nluc-β_2_AR thermostability in different detergent conditions. Data are pooled normalised values showing the mean ± standard deviation for the number of experimental replicates evident in the far-right graph (n≥2).

**Supplementary Figure 2:**
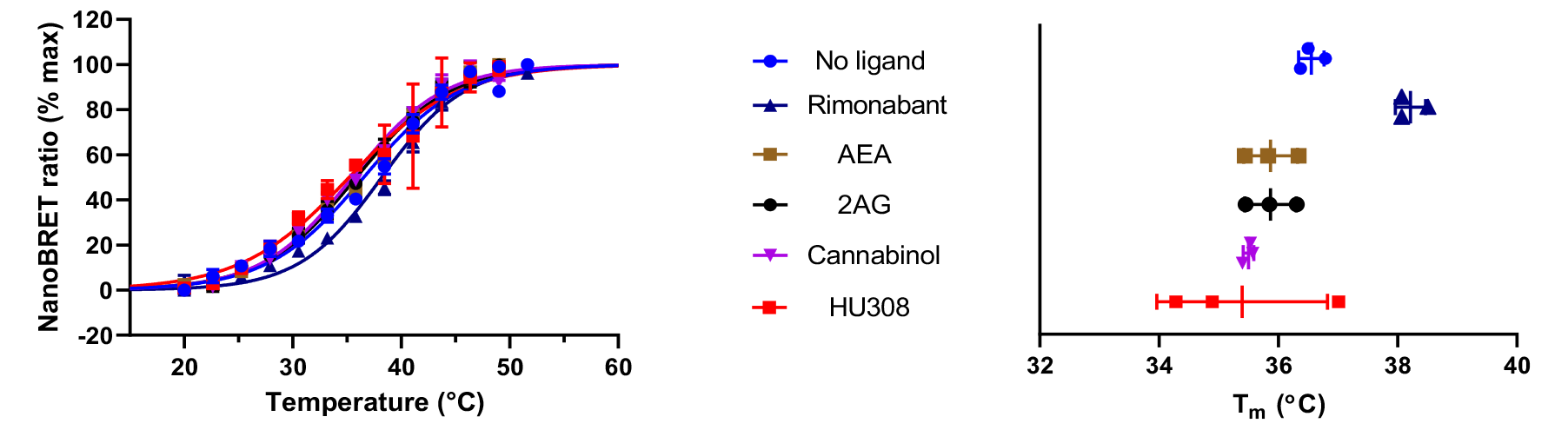
Thermostability of Nluc-CB_1_(91-472) solubilised in DDM/CHAPSO/CHS. Data are pooled normalised values showing the mean ± standard deviation for the number of experimental replicates evident in the far-right graph (n≥3). All ligands were present at a concentration of 20 µM.

**Supplementary Figure 3:**
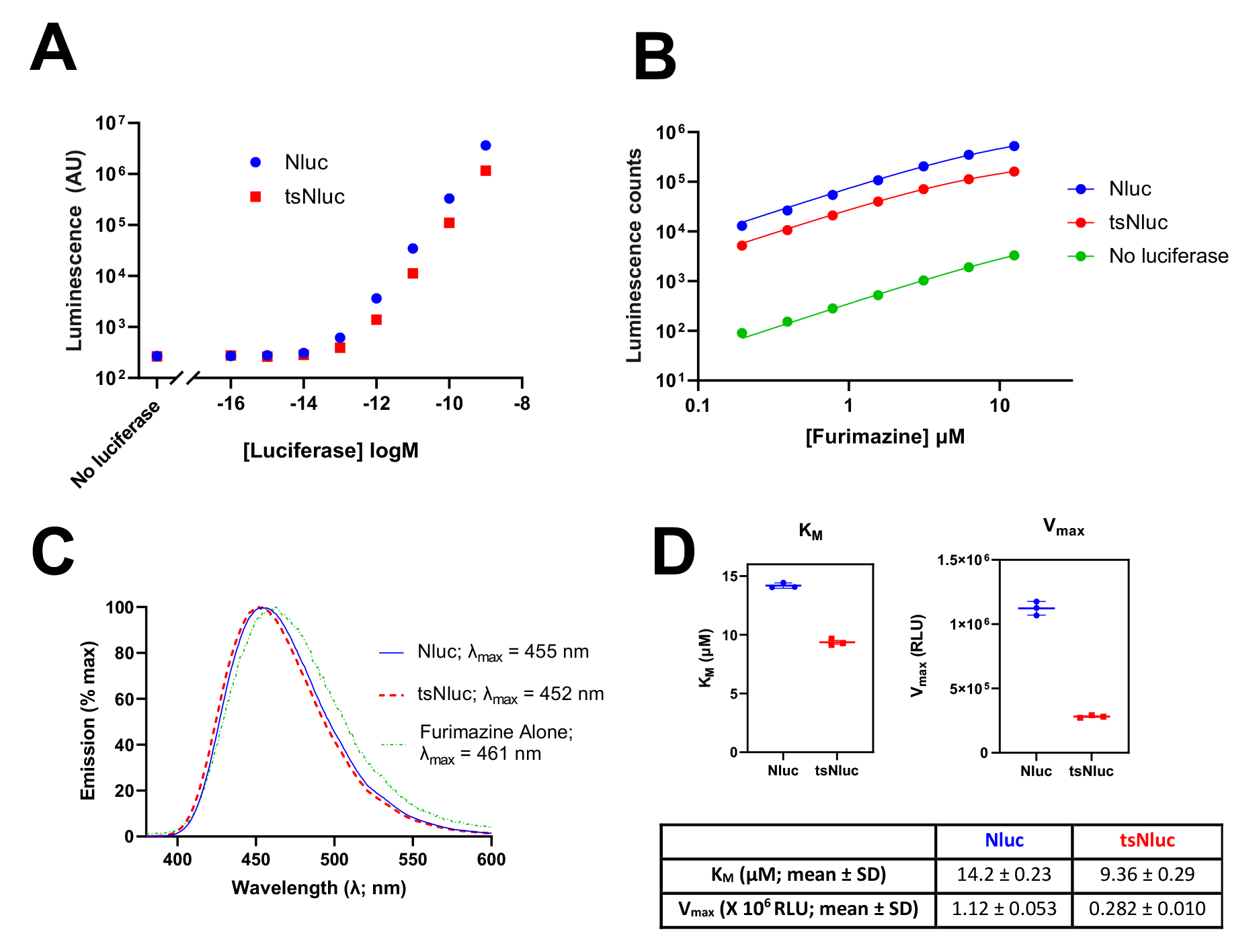
Characterisation of purified Nluc and tsNluc. **(A)** Different concentrations of luciferase proteins were mixed with 10 µM furimazine in a white 384-well optiplate and luminescence was measured with PHERAstar FSX plate reader with gain set to 2000 and 0.5 second measurement interval time. **(B)** Different concentrations of furimazine were mixed with 10 pM of purified luciferase and luminescence was measured with PHERAstar FSX plate reader with gain set to 3000 and 0.5 second measurement interval time. Data was fitted to a Michealis-Menten equation in GraphPad Prism to derive K_M_ and V_max_ values for Nluc and tsNluc. **(C)** Spectral scan of luminescence emission from 10 µM furimazine in the presence of absence of purified luciferases. **(D)** K_M_ and V_max_ determinations from **(B).** Data points and error bars represent the mean ± SD of 3 independent experiments performed in duplicate. Experiments were performed in CORE buffer at an ambient room temperature (26 - 27 ˚C at that time of year).

**Supplementary Figure 4:**
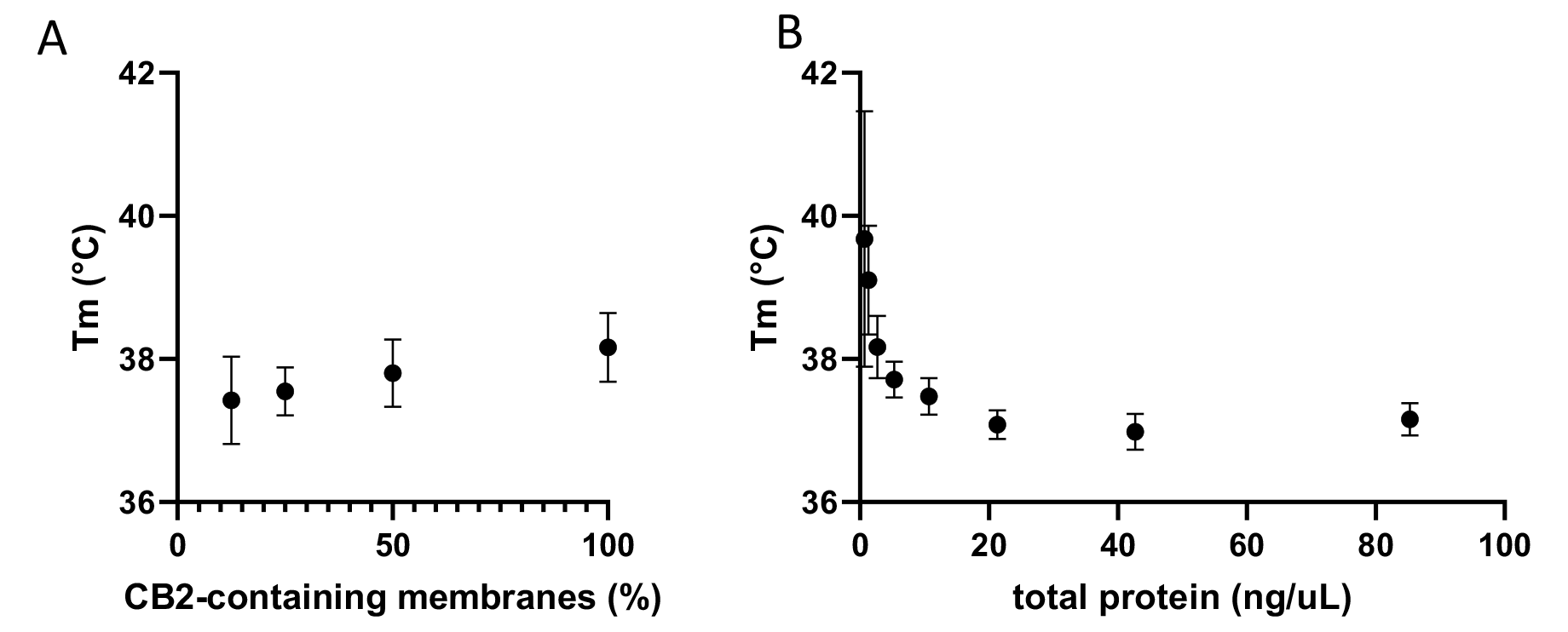
Dependence of T_m_ values on CB2 and membrane concentration (expressed as total protein). **(A)** Tm values for CB2-containing HEK293 membranes mixed in different ratios with “empty” non-transfected HEK293 membranes to a total of 26 ng/μL of protein. **(B)** Tm values for 0.67 ng/μL CB2- containing HEK293 membranes mixed with increasing concentrations of non-transfected HEK293 membranes. Samples were melted for 5 min at a temperature gradient 20-52°C in the presence of 0.1% DDM/0.05% CHAPSO/0.03% CHS. N=3, error bars are SEM.

**Supplementary Figure 5:**
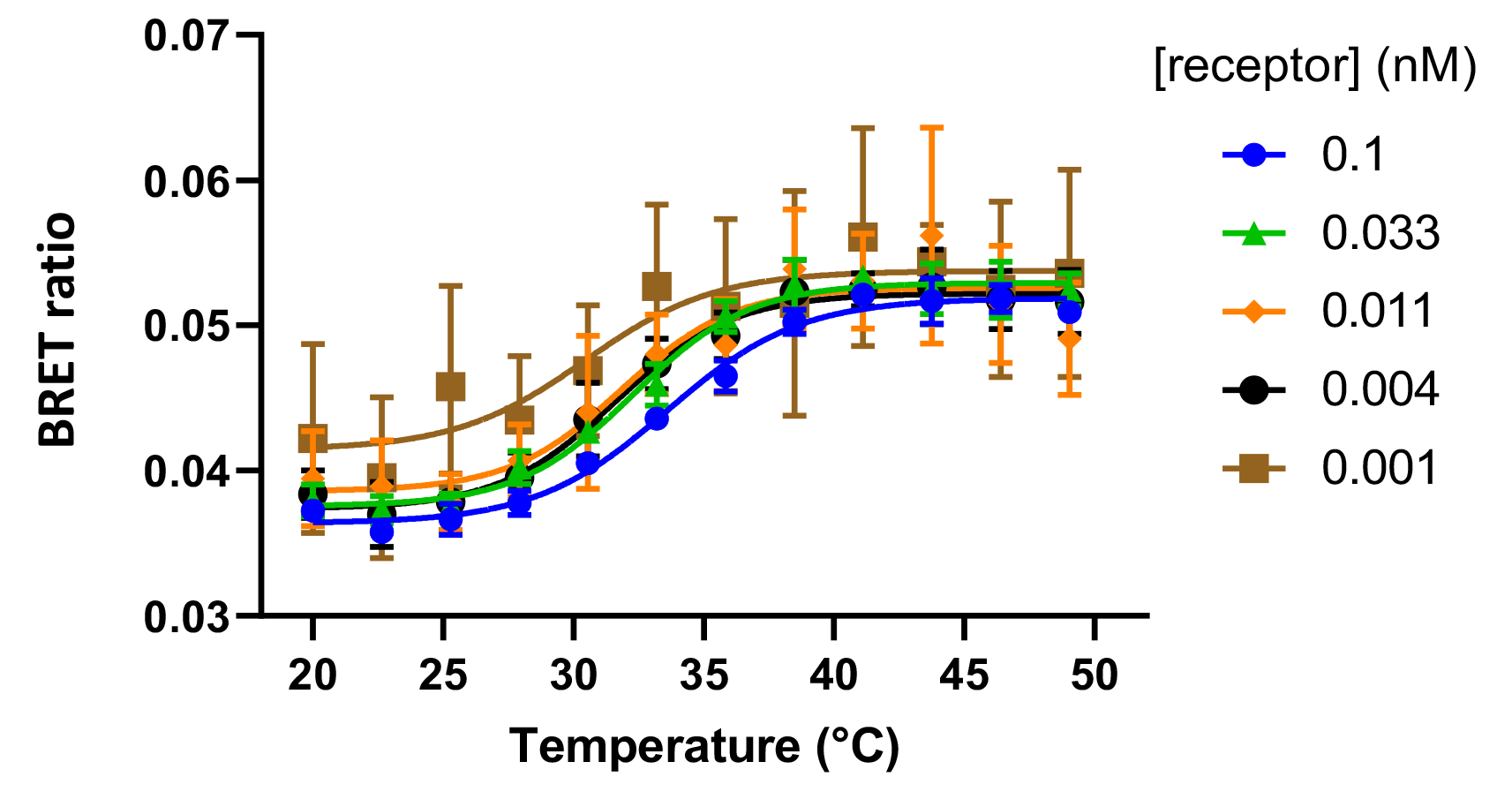
Assay sensitivity as determined by assessing the Thermostability of different concentrations of tsNluc-β_2_AR solubilised in DDM. Data are pooled normalised values showing the mean ± standard deviation. The different concentrations of receptor were estimated by diluting tsNluc and determining the level of luminescence at a fixed concentration of furimazine and comparing it to the levels of luminescence observed following dilution of the solubilised receptor.

**Supplementary Figure 6:**
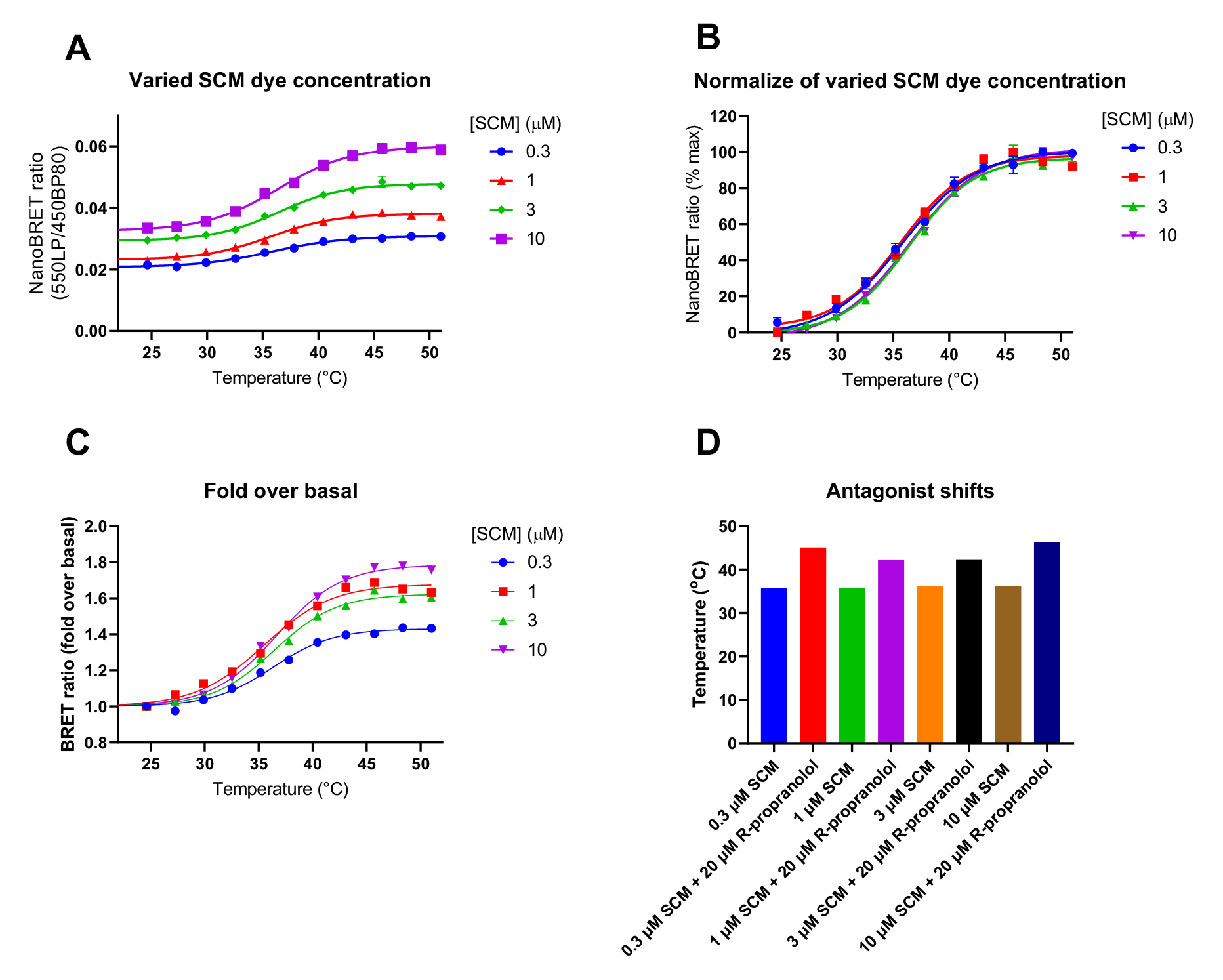
Effect of different SCM dye concentrations on *T*_m_ determinations from the tsNluc-β_2_AR solubilised in DDM. Increasing concentrations of SCM dye were incubated with a fixed level of tsNluc-β_2_AR and subject to a thermal gradient, *T*_m_ values were determined following the addition of a fixed concentration of furimazine (10 μM). BRET ratio values are shown in **(A)** and normalised data in **(B)** for the different dye concentrations. Fold over basal values are shown in **(C)**, and demonstrate the potential for signal improvement when increasing SCM concentration from 0.3 to 10 μM. The effect of dye concentration on the antagonist shift observed is shown in **(D)**. Data are single values determined from a single experiment.

## Supplementary Information 1: Amino acid sequences of expression constructs used in this study

**_>_ pcDNA4/TO SigPep-TwinStrep-Nluc-CB_2_**

MRLCIPQVLLALFLSMLTGPGEGSASDIGAPAFKSVQTGEFTAAAGSAWSHPQFEK GGGSGGGSGGSAWSHPQFEKGSGGSEDLMVFTLEDFVGDWRQTAGYNLDQVLE QGGVSSLFQNLGVSVTPIQRIVLSGENGLKIDIHVIIPYEGLSGDQMGQIEKIFKVVYP VDDHHFKVILHYGTLVIDGVTPNMIDYFGRPYEGIAVFDGKKITVTGTLWNGNKIIDE RLINPDGSLLFRVTINGVTGWRLCERILAPAGTMEECWVTEIANGSKDGLDSNPMK DYMILSGPQKTAVAVLCTLLGLLSALENVAVLYLILSSHQLRRKPSYLFIGSLAGADF LASVVFACSFVNFHVFHGVDSKAVFLLKIGSVTMTFTASVGSLLLTAIDRYLCLRYPP SYKALLTRGRALVTLGIMWVLSALVSYLPLMGWTCCPRPCSELFPLIPNDYLLSWLL FIAFLFSGIIYTYGHVLWKAHQHVASLSGHQDRQVPGMARMRLDVRLAKTLGLVLA VLLICWFPVLALMAHSLATTLSDQVKKAFAFCSMLCLINSMVNPVIYALRSGEIRSSA HHCLAHWKKCVRGLGSEAKEEAPRSSVTETEADGKITPWPDSRDLDLSDC*

**> pcDNA4/TO SigPep-TwinStrep-Nluc-β_2_AR**

MRLCIPQVLLALFLSMLTGPGEGSASDIGAPAFKSVQTGEFTAAAGSAWSHPQFEK GGGSGGGSGGSAWSHPQFEKGSGGSEDLMVFTLEDFVGDWRQTAGYNLDQVLE QGGVSSLFQNLGVSVTPIQRIVLSGENGLKIDIHVIIPYEGLSGDQMGQIEKIFKVVYP VDDHHFKVILHYGTLVIDGVTPNMIDYFGRPYEGIAVFDGKKITVTGTLWNGNKIIDE RLINPDGSLLFRVTINGVTGWRLCERILAPAGTMGQPGNGSAFLLAPNGSHAPDHD VTQQRDEVWVVGMGIVMSLIVLAIVFGNVLVITAIAKFERLQTVTNYFITSLACADLV MGLAVVPFGAAHILMKMWTFGNFWCEFWTSIDVLCVTASIETLCVIAVDRYFAITSP FKYQSLLTKNKARVIILMVWIVSGLTSFLPIQMHWYRATHQEAINCYANETCCDFFT NQAYAIASSIVSFYVPLVIMVFVYSRVFQEAKRQLQKIDKSEGRFHVQNLSQVEQDG RTGHGLRRSSKFCLKEHKALKTLGIIMGTFTLCWLPFFIVNIVHVIQDNLIRKEVYILLN WIGYVNSGFNPLIYCRSPDFRIAFQELLCLRRSSLKAYGNGYSSNGNTGEQSGYHV EQEKENKLLCEDLPGTEDFVGHQGTVPSDNIDSQGRNCSTNDSLL*

**> pcDNA4/TO SigPep-TwinStrep-Nluc-CB_1_(91-472)^#^**

MRLCIPQVLLALFLSMLTGPGEGSASDIGAPAFKSVQTGEFTAAAGSAWSHPQFEK GGGSGGGSGGSAWSHPQFEKGSGGSEDLMVFTLEDFVGDWRQTAGYNLDQVLE QGGVSSLFQNLGVSVTPIQRIVLSGENGLKIDIHVIIPYEGLSGDQMGQIEKIFKVVYP VDDHHFKVILHYGTLVIDGVTPNMIDYFGRPYEGIAVFDGKKITVTGTLWNGNKIIDE RLINPDGSLLFRVTINGVTGWRLCERILAPAGTENEENIQCGENFMDIECFMVLNPS QQLAIAVLSLTLGTFTVLENLLVLCVILHSRSLRCRPSYHFIGSLAVADLLGSVIFVYS FIDFHVFHRKDSRNVFLFKLGGVTASFTASVGSLFLTAIDRYISIHRPLAYKRIVTRPK AVVAFCLMWTIAIVIAVLPLLGWNCEKLQSVCSDIFPHIDETYLMFWIGVTSVLLLFIV YAYMYILWKAHSHAVRMIQRGTQKSIIIHTSEDGKVQVTRPDQARMDIRLAKTLVLIL VVLIICWGPLLAIMVYDVFGKMNKLIKTVFAFCSMLCLLNSTVNPIIYALRSKDLRHAF RSMFPSCEGTAQPLDNSMGDSDCLHKHANNAASVHRAAESCIKSTVKIAKVTMSV STDTSAEAL*

**> pcDNA4/TO SigPep-TwinStrep-tsNluc-CB_2_**

MRLCIPQVLLALFLSMLTGPGEGSASDIGAPAFKSVQTGEFTAAAGSAWSHPQFEKGGGSGGGSGGSAWSHPQFEKGSGGSEDLMVFTLEDFVGDWEQTAAYNLDQVLEQGGVSSLLQNLAVSVTPIQRIVRSGENALKIDIHVIIPYEGLSADQMAQIEEVFKVVYPVDDHHFKVILPYGTLVIDGVTPNMLNYFGRPYEGIAVFDGKKITVTGTLWNGNKIIDERLITPDGSMLFRVTINGVSGWRLFKKISPAGTMEECWVTEIANGSKDGLDSNPMKDYMILSGPQKTAVAVLCTLLGLLSALENVAVLYLILSSHQLRRKPSYLFIGSLAGADFLASVVFACSFVNFHVFHGVDSKAVFLLKIGSVTMTFTASVGSLLLTAIDRYLCLRYPPSYKALLTRGRALVTLGIMWVLSALVSYLPLMGWTCCPRPCSELFPLIPNDYLLSWLLFIAFLFSGIIYTYGHVLWKAHQHVASLSGHQDRQVPGMARMRLDVRLAKTLGLVLAVLLICWFPVLALMAHSLATTLSDQVKKAFAFCSMLCLINSMVNPVIYALRSGEIRSSAHHCLAHWKKCVRGLGSEAKEEAPRSSVTETEADGKITPWPDSRDLDLSDC*

**> pJ411 His-TEV-Nluc**

MKKHHHHHHHHHHENLYFQGGSVFTLEDFVGDWRQTAGYNLDQVLEQGGVSSLFQNLGVSVTPIQRIVLSGENGLKIDIHVIIPYEGLSGDQMGQIEKIFKVVYPVDDHHFKVILHYGTLVIDGVTPNMIDYFGRPYEGIAVFDGKKITVTGTLWNGNKIIDERLINPDGSLLFRVTINGVTGWRLCERILA*

**> pJ411 HIS-TEV-TsNluc** MKKHHHHHHHHHHENLYFQGGSVFTLEDFVGDWEQTAAYNLDQVLEQGGVSSLLQNLAVSVTPIQRIVRSGENALKIDIHVIIPYEGLSADQMAQIEEVFKVVYPVDDHHFKVILPYGTLVIDGVTPNMLNYFGRPYEGIAVFDGKKITVTGTLWNGNKIIDERLITPDGSMLFRVTINGVSGWRLFKKIS*

## Supplementary Information 2: Amino acid alignment of Nluc and tsNluc

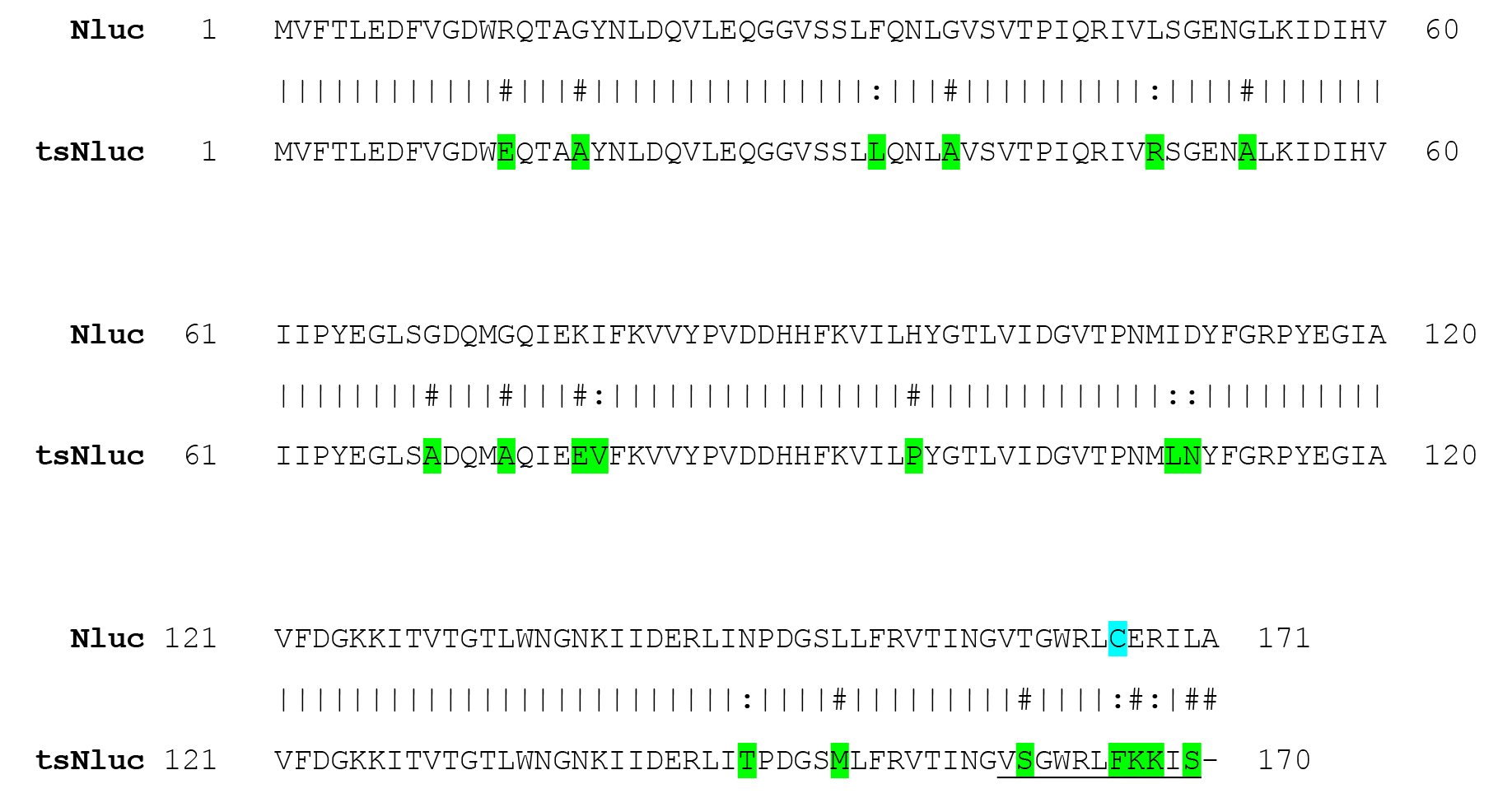

Amino acid alignment of Nluc and tsNluc. The split Nanoluciferase fragment sequences of LgBit and HiBiT (VSGWRLFKKIS) were joined to create tsNluc. Residues shaded green show mutations of tsNluc from the Nluc sequence. Note the C166F mutation which results in tsNluc containing no cysteine residues.

## Footnotes

\# The full-length CB_1_ receptor contains an unusually long (around 117 amino acids) and likely unstructured N-terminal domain. It was therefore truncated at the N-terminus in order to bring the Nluc tag in proximity with the transmembrane helices.

